# Aging shapes the population-mean and ‐dispersion of gene expression in human brains

**DOI:** 10.1101/039933

**Authors:** Candice L. Brinkmeyer-Langford, Jinting Guan, Guoli Ji, James J. Cai

## Abstract

Human aging is associated with cognitive decline and an increased risk of neurodegenerative disease. Our objective for this study was to evaluate potential relationships between age and variation in gene expression across different regions of the brain. We analyzed the Genotype-Tissue Expression (GTEx) data from 54 and 101 tissue samples across 13 brain regions in post-mortem donors of European descent aged between 20 and 70 years at death. After accounting for the effects of covariates and hidden confounding factors, we identified 1,446 protein-coding genes whose expression in one or more brain regions is correlated with chronological age at a false discovery rate of 5%. These genes are involved in various biological processes including apoptosis, mRNA splicing, amino acid biosynthesis, and neurotransmitter transport. The distribution of these genes among brain regions is uneven, suggesting variable regional responses to aging. We also found that the aging response of many genes, e.g., *TP37* and *C1QA*, depends on individuals’ genotypic backgrounds. Finally, using dispersion-specific analysis, we identified genes such as *IL7R*, *MS4A4E*, and *TERF1/TERF2* whose expressions are differentially dispersed by aging, i.e., variances differ between age groups. Our results demonstrate that age-related gene expression is brain region-specific, genotype-dependent, and associated with both mean and dispersion changes. Our findings provide a foundation for more sophisticated gene expression modeling in the studies of age-related neurodegenerative diseases.

## Introduction

Aging is a natural process bestowed upon those fortunate enough to participate in the journey, and the progression of age has profound impacts on physical and mental health. The mechanisms underlying age-related cognitive decline and increased risk of neurodegenerative disease remain unclear, though both decline and disease are universally common; therefore, it is critically important to understand the effects of aging on the human brain. One way to approach this goal is to detect the gene expression changes in the human brain during the aging process (Lu et al., 2004). Brain transcriptomic studies hold promise for better understanding the role of aging in both normal brain activity and the development of neurodegenerative disease. The advent of high-throughput sequencing has allowed the study of genome-wide patterns of change in gene expression associated with aging.

In recent years a number of studies on age-related gene expression have been published, such as (Glass et al., 2013;Peters et al., 2015;Sood et al., 2015;Yang et al., 2015), though some have either overlooked the central nervous system or focused on the brain *in toto* for comparisons with other organs and tissues. Meanwhile, mounting evidence shows that the human brain is functionally heterogeneous, with different sub-regions showing distinct functions, cell-type compositions, and gene expression patterns (Fraser et al., 2005;Oldham et al., 2008). In fact, the development and functions of separate anatomical regions of the brain are guided by specific, and often independent, networks of gene expression, e.g., (Kang et al., 2011;Miller et al., 2014;Tebbenkamp et al., 2014). Not surprisingly, therefore, different regions of the brain differ in their susceptibilities to diseases. For example, the hippocampus is affected in Alzheimer’s disease, while Parkinson’s disease affects the substantia nigra (Hyman et al., 1984;Jellinger, 1986). The striatum of the basal ganglia is a primary region affected in Huntington’s disease (Vonsattel et al., 1985). Neurodegenerative diseases tend to be unique to the affected brain sub-regions (Graveland et al., 1985;Fearnley and Lees, 1991;Francis et al., 1999;Neumann et al., 2006;Yao et al., 2015).

In the present study, we focused on the differential gene expression associated with age in multiple brain regions. Rather than evaluating only single genes, we also applied factor analysis (Anand Brown et al., 2015) to identify functional gene sets that are associated with age. More important, our analysis was directed to gene expression dispersion (e.g., variance in gene expression across samples or other measures of dispersion) as a metric to reveal a new model of age-related gene expression patterns. Several studies in humans and model organisms have suggested that age may influence the level of phenotypic dispersion of the population. Thus, to achieve a comprehensive picture of brain aging, we included the dispersion-specific analyses of gene expression with age.

## Methods

### GTEx brain tissues and expression data

The Genotype-Tissue Expression (GTEx) project was established to determine how genetic variation affects normal gene expression in human tissues, ultimately to inform the study of human diseases (GTEx_Consortium, 2015). The project has collected multiple different human tissues from each of hundreds of donors to isolate nucleic acids from the tissues and perform genotyping, gene expression profiling, whole genome sequencing, and RNA sequencing (RNA-seq) analyses. Among these many tissues, there is a plethora of samples from a handful of sub-regions of the brain, from which expression data sets were generated for the GTEx project and used in the present study.

The expression data (v6, October 2015 release) for brain specimens of post-mortem donors were obtained from the GTEx portal website (http://www.gtexportal.org/). The data was generated using RNA-seq with tissues initially sampled from two brain regions: cerebellum and cortex, preserved using the PAXgene tissue preservation system (Groelz et al., 2013), and with tissues subsequently sampled from frozen brains in following regions: amygdala, anterior cingulate cortex (BA24), caudate (basal ganglia), cerebellar hemisphere, frontal cortex (BA9), hippocampus, hypothalamus, nucleus accumbens (basal ganglia), putamen (basal ganglia), spinal cord (cervical c-1), and substantia nigra (Carithers et al., 2015;GTEx_Consortium, 2015). From the downloaded data, we extracted the whole-gene level RPKM (Reads Per Kilobase of transcript per Million mapped reads) values for protein-coding genes. The data for different brain regions was quantile normalized, and log2 transformed, separately. For each region, 10% lowly expressed genes were excluded from data analysis based on their mean expression level across samples. The donor’s information of gender, body mass index (BMI), and sample’s ischemic time were also downloaded. Tissue samples from donors of non-European ancestry were excluded from the subsequent data analyses. The number of remaining samples of 13 brain regions ranged between 54 and 101 (**Table 1**).

**Table 1.**
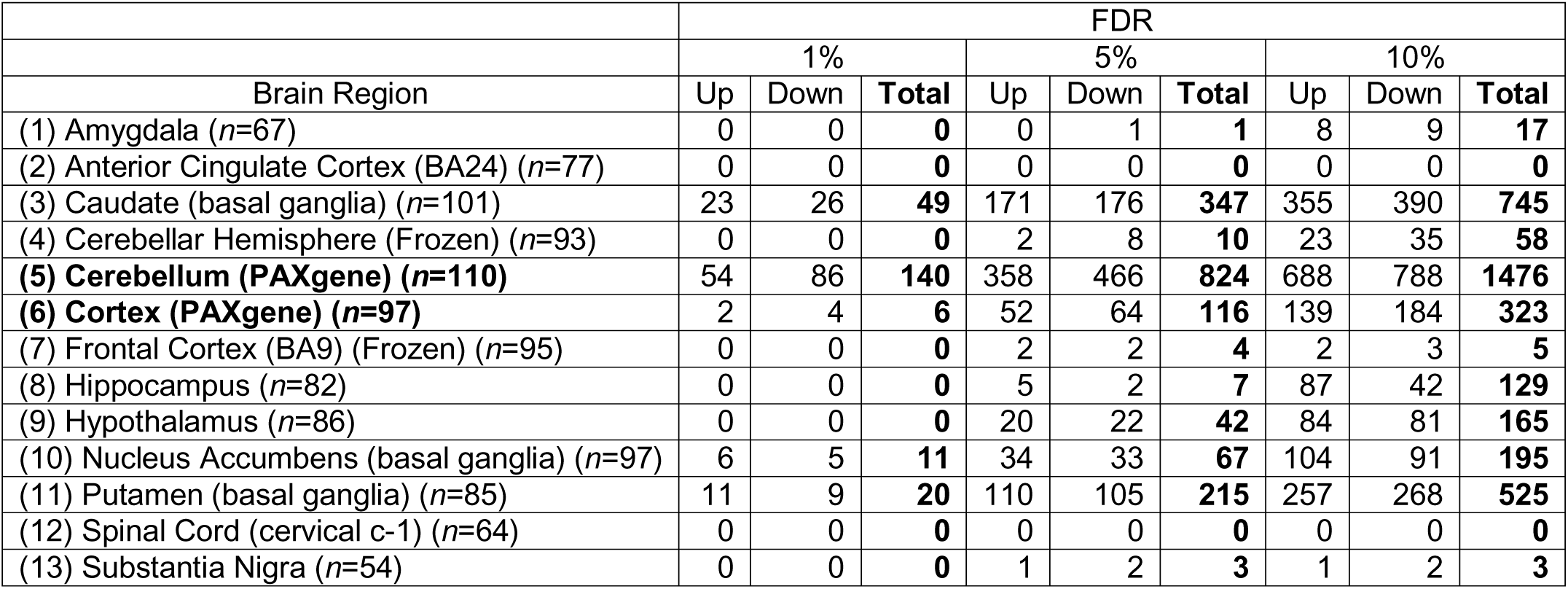
Numbers of age-related genes in 13 GTEx brain regions. Columns “Up” and “Down” list the numbers of up-regulated and down-regulated aging genes, respectively. Results derived from using three different FDR cutoffs (1%, 5%, and 10%) are shown. Two regions, (5) Cerebellum (PAXgene) and (6) Cortex (PAXgene), for which samples were preserved using the PAXgene tissue preservation system, are highlighted in bold.

### Accounting for confounding factors using PEER algorithm

Prior to the regression analysis, we used a two-step approach based on the PEER algorithm (Stegle et al., 2012) to control for known covariates as well as hidden data structures in the GTEx expression data. For each region, PEER was first used to discover patterns of common variation across the entire data set and create up to 15 assumed global hidden factors. In doing so, the known covariates, including the donors’ age, gender and BMI for all samples from the 13 regions, were included in the PEER models. Also, for samples from (5) cerebellum (PAXgene) and (6) cortex (PAXgene), ischemic time was included as one additional covariate. Note that, at this step, the age of donors was included to enable the PEER to discover correlated patterns across global structured data (O. Stegle, personal communication, November 11, 2015). Next, the correlation between each of the 15 constructed factors and age was tested with a data set of each region. The factor(s) showing a Pearson’s correlation test *P*-value smaller than 0.05 were excluded. The remaining factors (denoted *PC*_*k*_, where 1 □≤□*k*□≤□*N* and *N* is the number of factors), along with non-age covariates (i.e., known covariates excluding age), were used as a new set of covariates in the regression analysis. Furthermore, in the pathway-based factor analysis (described below), the remaining factors and non-age covariates were supplied to PEER as a new set of covariates and were regressed out. In this way, the effects of all known covariates other than age and hidden data structures, which could potentially confound the subsequent analyses, were removed. The residual values of the regression were used as a new, corrected gene expression data in the subsequent analyses.

### Analysis of gene expression with age using linear regression model

For each region, we modeled gene expression using the following linear regression model:

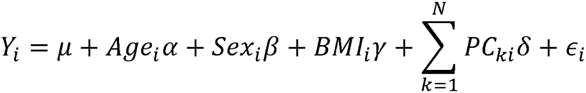

where *Y*_*i*_ is the expression level of a given gene in sample *i*; *Age*_*i*_, *Sex* and *BMI*_*i*_ are the age, sex and BMI of sample *i* with regression coefficients *αβ* and *γ* respectively; *PC*_*ki*_ (1□≤□*k*□≤□*N*) is the value of the *k*-th hidden factors for the *i*-th sample with regression coefficient *δ*; *N* is the total number of factors uncorrelated with age; □_*i*_, is the error term, and *μ* is the regression intercept.

We fitted the model in Matlab with the `fitlm` function in the Statistics toolbox. For each gene, a least square approach was used to estimate the regression coefficients. If *α* was significantly deviated from 0, the gene was considered to be age-associated. A gene was considered up-regulated with age if *α* □>□0 and down-regulated if *α* □<0□0.

Throughout this study, the GO term enrichment analysis was carried out using the DAVID Bioinformatics Resource server (Dennis et al., 2003). The FDR adjustment on the *P*-values was made using the Benjamini–Hochberg procedure (Benjamini and Hochberg, 1995).

### Pathway-based factor analysis of gene expression associated with age

The rationale behind pathway-based factor analysis is that a statistical factor analysis (e.g., PEER) can not only remove noise components from high-dimensional data but also derive factors summarizing pathway expression to analyze the relationships between expression and aging (Anand Brown et al., 2015). We used the pathway-based factor analysis to analyze the correlation between age and gene expression of GO-term defined gene sets. We first applied PEER to the whole gene expression matrix for each brain region to regress out global factors. The residual expression levels were treated as new expression data sets; for a given GO-term gene set, PEER was used to construct factors. The constructed factors on the gene sets were taken as concise summaries of common expression variation across each set. These factor values were considered as phenotypes and referred to as phenotype factors. Subsequently, by looking for associations between these new phenotype factors and age, we discovered groups of functionally related genes with a common response to aging.

### Detecting effect of genotype-by-age interaction on gene expression

To investigate the genotype-by-age interaction contribution to gene expression, we included the genotype-by-age interaction term to the linear regression model described above. As a contributing factor to the gene expression variance, the significance of this interaction term was assessed for each gene after the model was fitted. Sample donors’ genotype data was downloaded from dbGaP under accession number phs000424.v6.p1 (October 2015). At each polymorphic site, an individual’s genotype was denoted with 0, 1, or 2 based on the number of non-reference alleles, respectively. SNPs with minor allele frequency greater than 15% were included for the test. To keep the overall computing time feasible, we randomly selected 2,000 genes genome-wide and also included only SNPs with minor allele frequency greater than 15% in the analysis, which was run on the high-performance computing cluster of Texas A&M Institute for Genome Sciences and Society (TIGSS).

### Test for expression heteroscedasticity between age groups

To compare the level of gene expression dispersion between age groups, we used Levene’s tests. The test examines if the gene expression levels of different age groups have equal deviations from the group means. Let *x*_*kj*_ be a set of *j*=*1*,…,*n*_*k*_ observations in each of *k*=*1*,…,*g* age groups. Levene’s test statistic is the ANOVA F-ratio comparing the *g* groups, calculated on the absolute deviations *z*_*kj*_ = |*x*_*kj*_ – *x*̅_*k*_.|, where 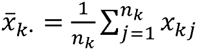 is the group means. To extend the analytical framework to muliple genes, we used the Mahalanobis distance (MD)-based generalization of Levene’s test (Anderson, 2006). A robust version of MD was used to quantify the distance from individual sample *i* to the multivariate centroid of all samples: 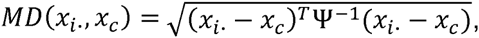, where *x*_*i*_ is the vector of expression of genes in sample *i*; *x*_*c*_ is the location estimator based on the minimum covariance determinant (Rousseeuw and Van Driessen, 1999); and ψ is the scattering estimator. Let *MD*_*kj*_ be a set of *j*=*1*,…,*n*_*k*_ observations in each of *k*=*1*,…,*g* age groups, Levene’s test literally performs ANOVA on *MD*_*kj*_, given the absolute deviation 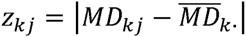 with the group means 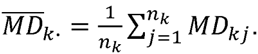.

## Results

### Identification of brain region-specific age-related genes

The data of transcriptomic profiles for brain tissues of 169 European ancestry donors aged 20 – 70 years at death (GTEx_Consortium, 2015) was downloaded from the GTEx portal website.The tissue samples were collected from 13 regions (or subareas) of the human brain, namely (1) amygdala, (2) anterior cingulate cortex [Brodmann area 24 (BA24)], (3) caudate (basal ganglia), (4) cerebellar hemisphere, (5) cerebellum (PAXgene), (6) cortex (PAXgene), (7) frontal cortex (BA9), (8) hippocampus, (9) hypothalamus, (10) nucleus accumbens (basal ganglia), (11) putamen (basal ganglia), (12) spinal cord (cervical c-1), and (13) substantia nigra. We excluded samples derived from non-European donors. The final data matrices, including 54 – 101 samples all from European donors, were normalized separately by different brain regions (**Methods**).

We used linear regression models, controlling for covariates and hidden confounding factors (**Methods**), to identify genes whose expression is correlated or anti-correlated with chronological age. **Table 1** shows the numbers of age-related genes in the 13 brain regions at the false discovery rate (FDR) of 1%, 5%, and 10%. The number of genes identified in most regions increases with the relaxation of FDR cutoff except in (2) anterior cingulate cortex and (12) spinal cord where no genes were identified. At FDR of 5%, 1,446 distinct age-related genes across all regions were identified (**Supplementary Table S1**). Of these, 155 were found in more than one region of the brain: seven were identified in four, 21 in three, and 127 in two brain regions. For each of these "multi-hit" genes, the directions of expression response to aging were the same in the different brain regions where the gene was identified. For comparison, in a previous study, using microarray data from ten regions of 100 post-mortem brains aged from 16 to 83 years, Glass et al. (2013) identified 14 age-related genes. Out of the 14 genes, six (*HSD11B1, MS4A6A, MT1G, PTPN3, SLC7A5*, and *WWC2*) are among our 5% FDR age-related genes, showing consistent directions of expression response to aging. In another previous study, Lu et al. (2004) compared frontal cortical samples from young and old adult individuals and identified 416 age-related genes whose expression differs by at least 1.5-fold. Out of the 416 genes, 61 (14.7%) are among our 5% FDR age-related genes.

**Table 1** also shows that some regions demonstrate significantly more age-related genes than others, suggesting distinct brain regions might have different levels of sensitivity or responsiveness to aging. To show that such regional specificities are not completely due to the difference in the number of samples from different brain regions used in the analysis, we repeated the identification of age-related genes by randomly subsampling samples to 54 (the minimal sample size) for all regions. We found that the number of identified genes decreased substantially as the sample size decreases, but the differences in numbers of identified genes between regions largely remained (**Supplementary Table S2**).

The numbers of age-related genes identified show a significant discrepancy between (4) cerebellar hemisphere and (5) cerebellum (PAXgene), which is unexpected because tissue samples of (4) and (5) were essentially from the same region of the cerebellum. Similarly, it is unexpected to see a great discrepancy in the numbers of age-related genes identified between (6) cortex (PAXgene) and (7) frontal cortex (BA9), because (6) and (7) were sampled from the same region of cortex. Indeed, clustering analysis based on the Euclidean distance between gene expression profiles confirmed that (4) and (5), as well as (6) and (7), are more similar to each other, respectively, than to other brain regions (**Supplementary Fig. S1**). We consider that the markedly fewer genes identified in (4) than (5), and in (7) than (6), may be attributed to whether or not the samples were subject to frozen storage before RNA-seq was performed. Among all GTEx brain specimens, only (5) and (6) were initially sampled "on site" from the post-mortem donors, while the rest were subsequently resampled after the brains were frozen and stored (Carithers et al., 2015;GTEx_Consortium, 2015). Thus, it is likely that the frozen-thaw cycle introduced extra expression variability to the samples [e.g., in (4) and (7)], resulting in the identification of fewer genes. To illustrate this further, we sought to examine the cross-region correlation between genes’ responsiveness to aging. We used each gene’s P-value against age in the linear regression model as the measure of the gene’s responsiveness to aging. We ranked genes by their P-values and then compared the ranks of genes across regions. If the correlations between (4) and (5) and between (6) and (7) were high, then we considered that the discrepancies in age-related gene numbers between (4) and (5), and between (6) and (7), were simply due to the effect of freezing on the statistical power of age-related gene detection, rather than on the gene expression regulation. **Fig. 1** shows the correlation matrix with Spearman correlation coefficient (SCC) between regions. Firstly, we found that the correlation (i.e., the similarity in gene rank) between anatomically closely related regions is higher. For example, the SCC between (3) caudate and (11) putamen, which both belong to basal ganglia, is the highest among all region pairs. Intriguingly, the second highest is between (4) and (5), despite of the considerable discrepancy in identified genes between the frozen and unfrozen cerebellar samples, reinforcing the point that samples from the same brain region are indeed more similar to each other with respect to the genes’ responsiveness to aging. Likewise, (6), (7) and (2) anterior cingulate cortex are more correlated with each other than with other regions. Thus, the human brain appears to have different aging patterns in the cerebellum, cortex, and basal ganglia [including (3) caudate, (10) nucleus accumbens, and (11) putamen] (**Fig. 1**). These results are consistent with the findings of a previous microarray-based gene expression study (Fraser et al., 2005).

**Fig. 1.**
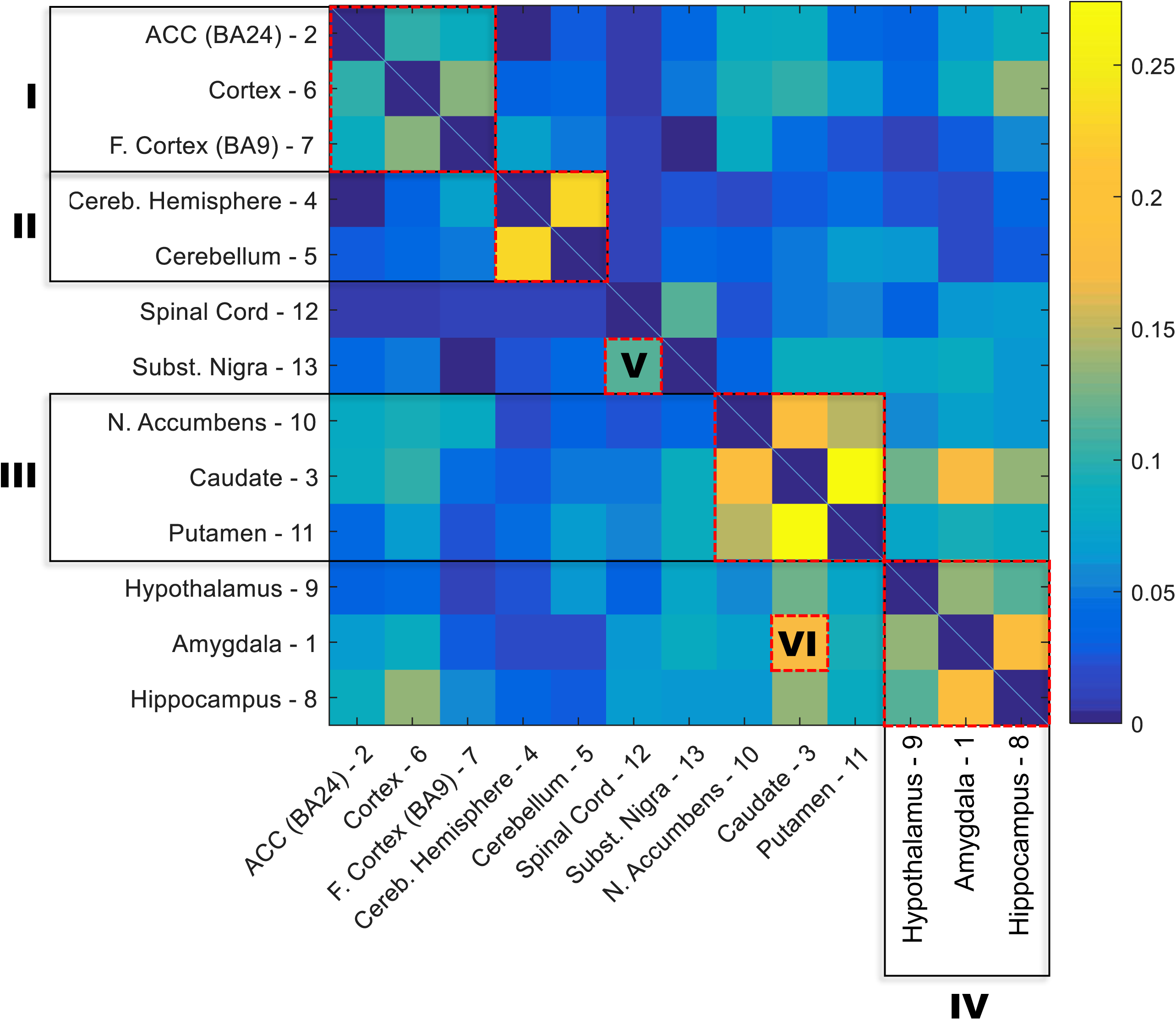
Correlation matrix for the responsiveness of genes to aging between 13 GTEx brain regions. The responsiveness to aging of a gene was measured with the P-value for slope coefficient in the linear regression model between age and gene expression. The correlation between each two brain regions was estimated with the nonparametric Spearman rank correlation coefficient between P-values of all genes in the two regions. The order of regions in the matrix was rearranged based on the similarity between regions. Six clusters of highly correlated regions are highlighted with red boxes: I – cortex; II – cerebellum; III – basal ganglia;IV – hypothalamus, amygdala, and hippocampus; V – substantia nigra and spinal cord; and IV – amygdala and caudate.

### Analysis of gene expression pathway factors associated with age

We set out to detect gene sets, in addition to single genes, with expression associated with age. We adopted the pathway-based factor analysis (Anand Brown et al., 2015) and applied it to 14,825 functional gene sets defined by gene ontology (GO) terms (**Methods**). As a result, 239 highly significant gene sets across the 13 brain regions were identified (*P* < 0.05, corrected using Bonferroni procedure for the total number of tested gene sets) (**Supplementary Table S3**). The related GO terms included: *neurogenesis* (GO:0022008), *neuron projection* (GO:0043005), *memory* (GO:0007613), and *regulation of synaptic plasticity* (GO:0048167). To obtain a broader functional overview of gene sets, we used the clustering method implemented in REVIGO (Supek et al., 2011) to summarize as many as 5,787 GO terms associated with age-related genes at 5% FDR significance. With REVIGO, these GO terms were evaluated against each other and clustered based on their context similarity. The TreeMap plots for the clusters were then generated, showing that the function of age-related gene sets points to a large collection of biological processes (BP) (**Fig. 2**) and molecular functions (MF) (**Supplementary Fig. S2**). For example, the top-level BP GO term clusters are represented by the terms apoptotic signaling pathway, aging, spliceosomal complex assembly, glutathione derivative biosynthesis, neurotransmitter transport, vitamin metabolism, reactive oxygen species metabolism, methylation, establishment or maintenance of cell polarity, and viral process (**Fig. 2**). The top-level MF GO term are listed in **Supplementary Fig. S2**. The largest clusters represented include growth factor binding, peptidase activity, phosphotransferase activity(alcohol group as acceptor), copper ion binding, symporter activity, heme binding, virus receptor activity, poly(A) RNA binding, and beta-amyloid binding (**Supplementary Fig. S2**).

**Fig. 2.**
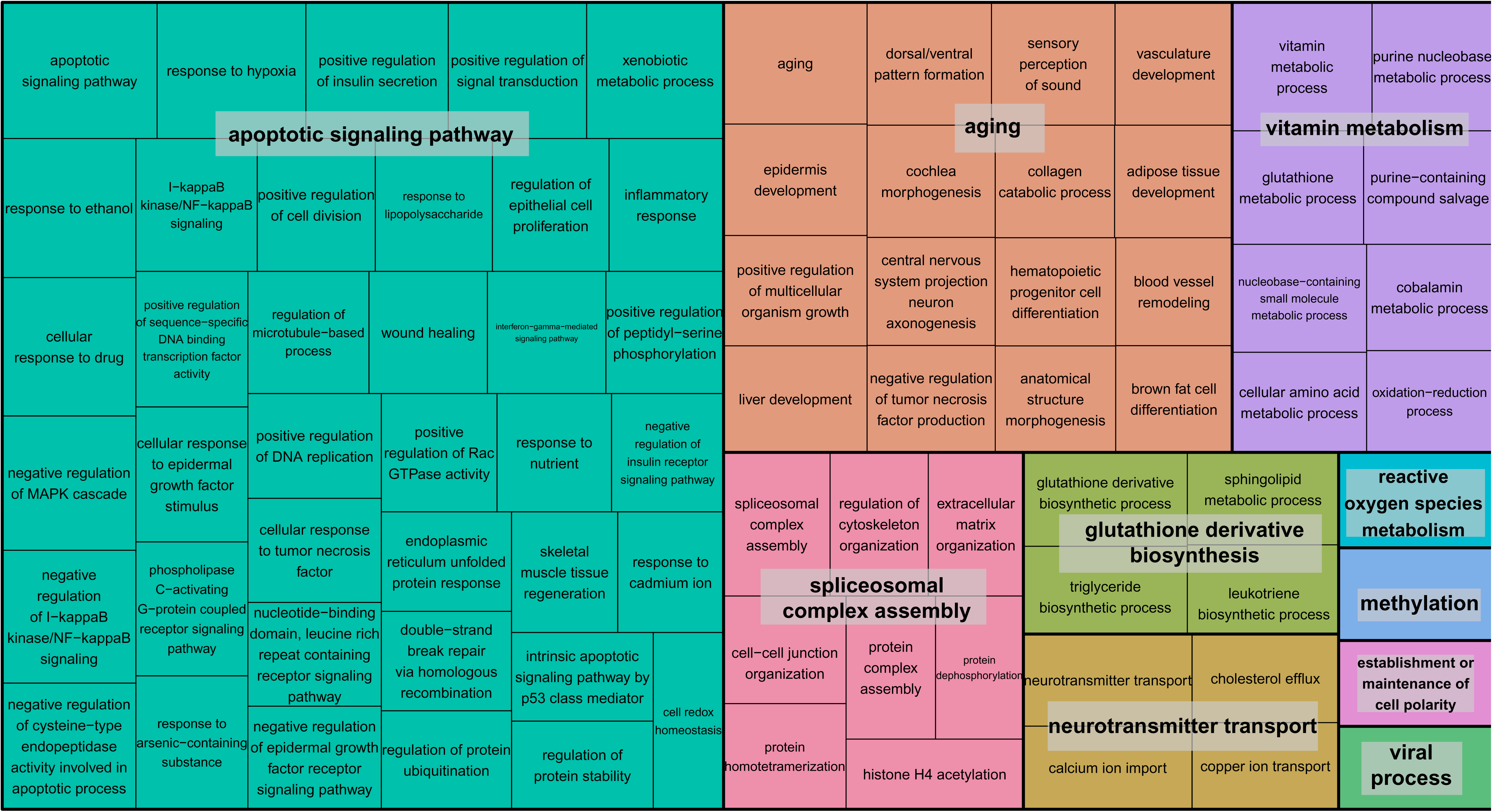
TreeMap view of GO-term clusters for age-related genes in the brain. The TreeMap was generated by using REVIGO with the input of all GO BP terms of age-related genes detected in 13 GTEx brain regions at the FDR of 5%. Each rectangle represents a single GO-term cluster. The size of the rectangles is proportional to the frequency of the GO term in the associated GO annotation database. The cluster representatives were joined into ‘superclusters’ of loosely related terms, visualized with different colors.

### Genotype-by-age interactions

We evaluated the interactions between genotype and age to assess how genetic background influences gene expression in the brain at different ages (**Methods**). These interactions could be used to develop strategies to predict individuals’ risks for specific conditions based on the associations between each individual’s genotype and expression changes anticipated for genes of interest. Other studies have identified age-related eQTL such as those connected with longevity (Walter et al., 2011;Erikson et al., 2016), AD [(Proitsi et al., 2014;Zhu et al., 2014); see also (Guerreiro et al., 2010) though this study did not identify any significant eQTL], and neurological conditions such as PD (Hernandez et al., 2012) and hippocampal sclerosis of aging (Nelson et al., 2015). In this study, we detected a number of interactions with high significance (nominal *P* < 10^−5^). A comprehensive list of SNPs and genes is provided (**Supplementary Table S4**), albeit none of these interactions survived multiple testing corrections due to the sheer large number of tests performed. We found it intriguing that certain genotypes seem to be more susceptible to the effects of aging on the expression of functionally significant genes. For example, genotypes of SNP rs55675298 can have different effects on expression of tumor protein p73 gene, *TP73* (**Fig. 3A**). For individuals with the GG genotype, there is an age-associated increase in *TP73* expression; individuals with GT or TT genotypes do not experience this increase, which could have profound health implications based on the potential roles of TP73 in conditions related to aging. TP73 is a member of the p53 transcription factor family and is located in a region that is frequently deleted in tumors, particularly neuroblastomas. Furthermore, *TP73* has been found to be critical for normal neuronal development and survival, making it a potential candidate gene for susceptibility to Alzheimer’s disease (AD) (Pozniak et al., 2000;Yang et al., 2000;Pozniak et al., 2002;Li et al., 2004;Wetzel et al., 2008). Another example of a relationship between SNP genotype and age-related gene expression involves *C1QA* and the SNP rs72788737 (**Fig. 3B**). Here again, the GG genotype seems to confer increased expression with age while GT/TT genotypes are not correlated with increased expression. Normal aging is associated with an increase in C1q protein (encoded by *C1QA)*, particularly in certain regions of the brain that are especially prone to degenerative diseases related to aging (Stephan et al., 2013). C1q contributes to an aging-related decrease in the regenerative capacity of certain tissues (Naito et al., 2012). Less C1q, on the other hand, may confer some protection against synapse loss and aging-related dysfunction of the hippocampus, e.g., (Stephan et al., 2013;Hong et al., 2016).

**Fig. 3.**
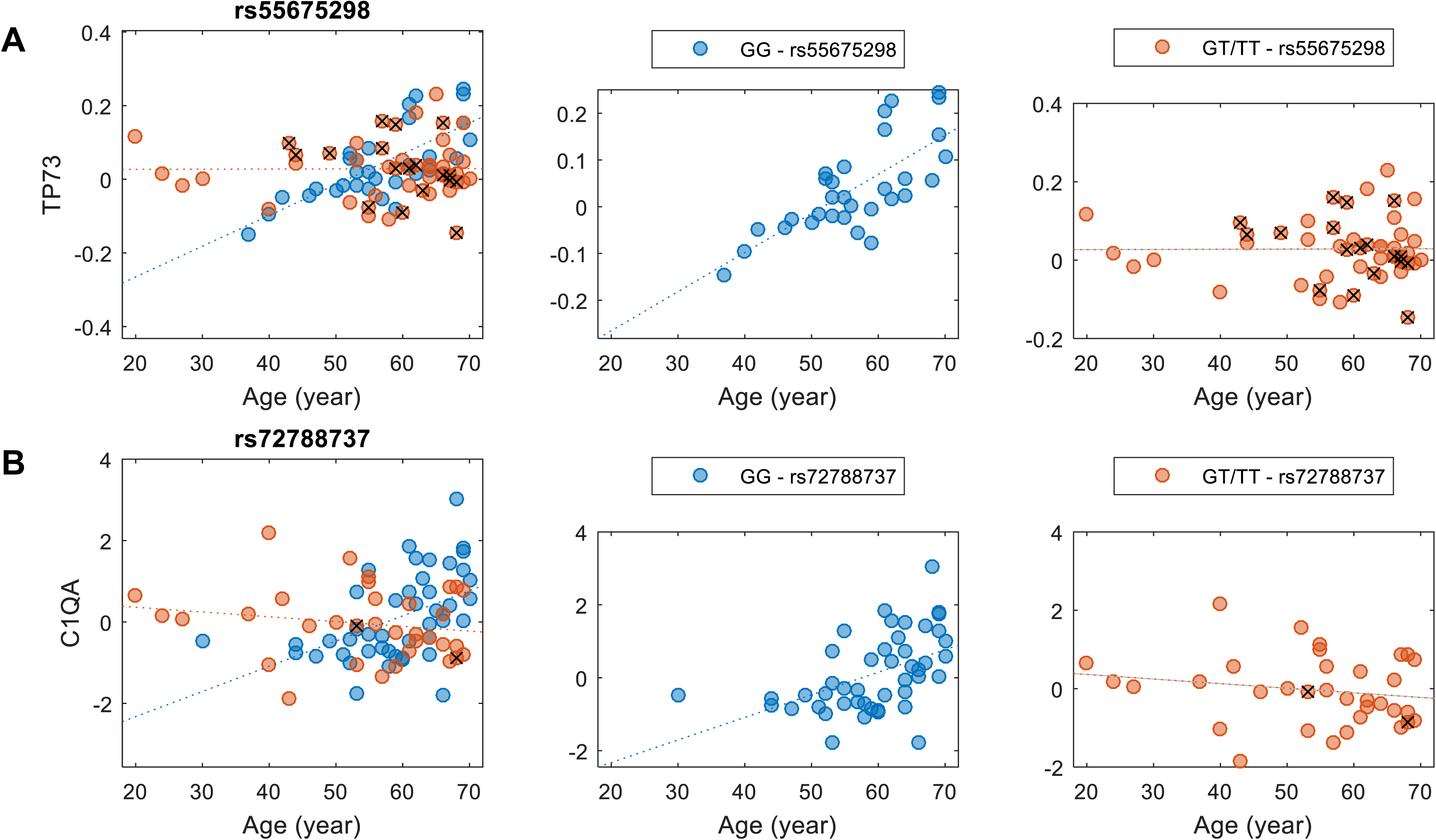
Examples of genotype-by-age interaction affecting the expression level of the gene. (**A**) The interaction between rs55675298 and age affecting *TP73* gene expression. (**B**) The interaction between rs72788737 and age affecting *C1QA* gene expression. For each subplot, the left panel shows all samples, the middle panel shows the major allele homozygous samples, and the right panel shows heterozygous and minor allele homozygous (with a cross) samples.

### Aging affects the population-level dispersion of gene expression in brain

Next, we focused on the identification of differentially variable (DV) genes. These genes show differences in the degree of dispersion in expression levels between age groups. Using Levene’s test, we identified 970 DV genes across brain regions (including 848 distinct genes) showing a significant difference in expression variance between young (20–60 years) and old (61–70 years) individuals at the significance level of FDR of 5% (**Supplementary Table S5**). About half of these genes show an increased expression variance in the old while the other half show a decreased variance. Across brain regions, the distribution of DV genes is more balanced (**Supplementary Table S6**), compared to the distribution of age-related genes. The region (6) cortex contains 108 DV genes, which is the most; while the (12) spinal cord contains as little as 40, which is the least. The top GO-term clusters for these DV genes include those related to sensory perception, peptide receptor activity, chemotaxis, peptidase inhibitor activity, and neurotransmitter binding (**Table 2**). **Fig. 4A, B** show two examples of DV genes—*IL7R* and *MS4A4E.* The expression dispersion of *IL7R* in the hippocampus is more pronounced in old than young adults (**Fig. 4A**). This gene is known for its possible role as a determinant of the rate of aging (Passtoors et al., 2015). In the other example, the expression dispersion of *MS4A4E* in the hippocampus also increases with age (**Fig. 4B**). This gene, as a member of the membrane-spanning four domains subfamily A gene cluster, plays a role in embryogenesis, oncogenesis, and the development of AD (Liang et al., 2001;Karagiannis et al., 2003;Hollingworth et al., 2011;Naj et al., 2011).

**Fig. 4.**
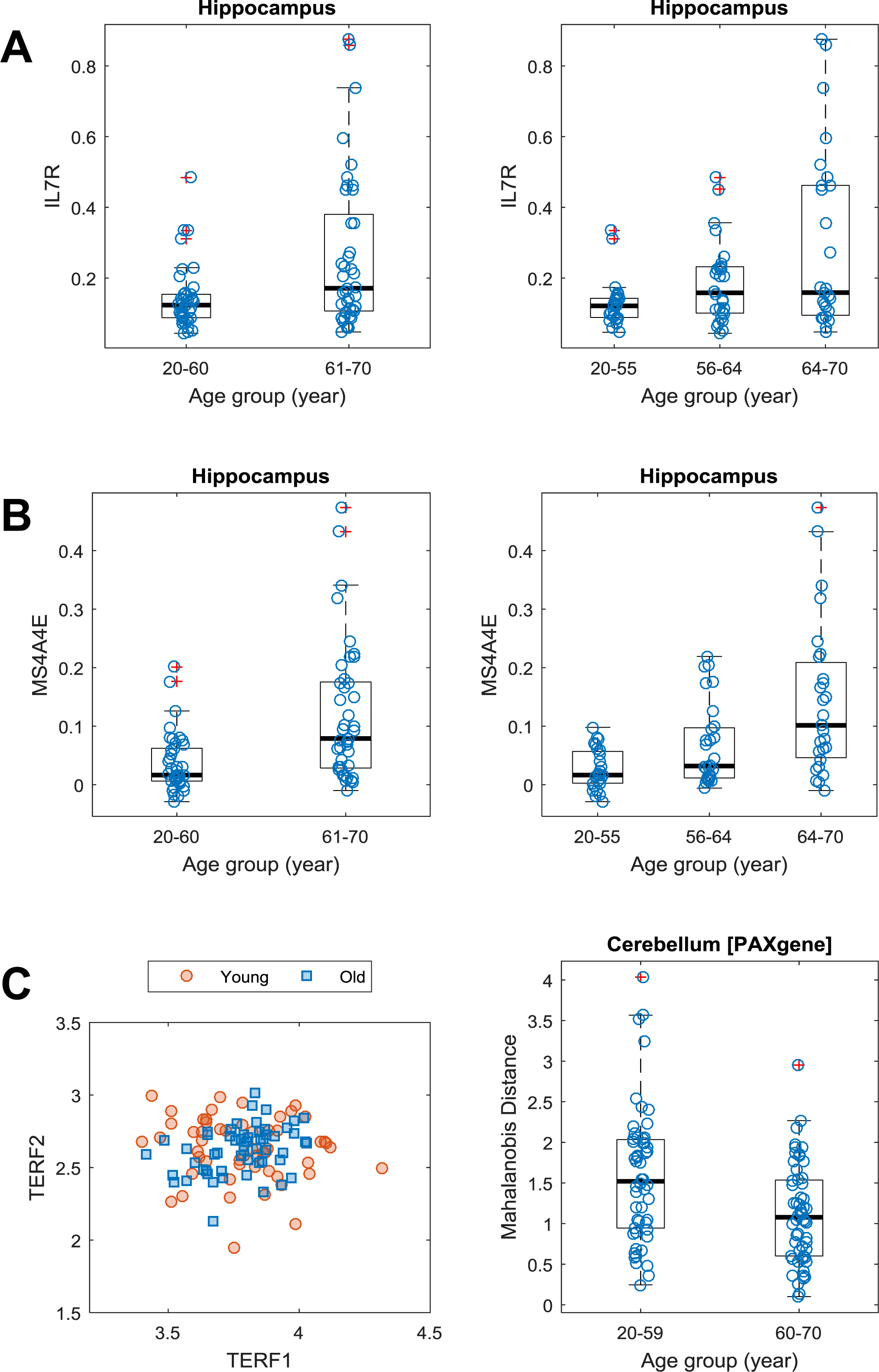
Differential gene expression dispersion between age groups. (**A**) Increased gene expression variance of *IL7R* in hippocampus between age groups (left: 20-60 years vs. 61-70 years; right: 20-55, 56-64, and 64-70 years). Each age group was plotted with jitter along the x-axis to show samples within each genotype. (**B**) Same as (**A**) but for *MS4A4E.* (**C**) Scatter plot (left) of expression levels of *TERF1* and *TERF2* with data points grouped by young (20-59 years) and old (60-70 years) ages. Boxplot (right) of MDs of young and old samples’ *TERF1* and *TERF2* expression profiles to the population center.

**Table 2.**
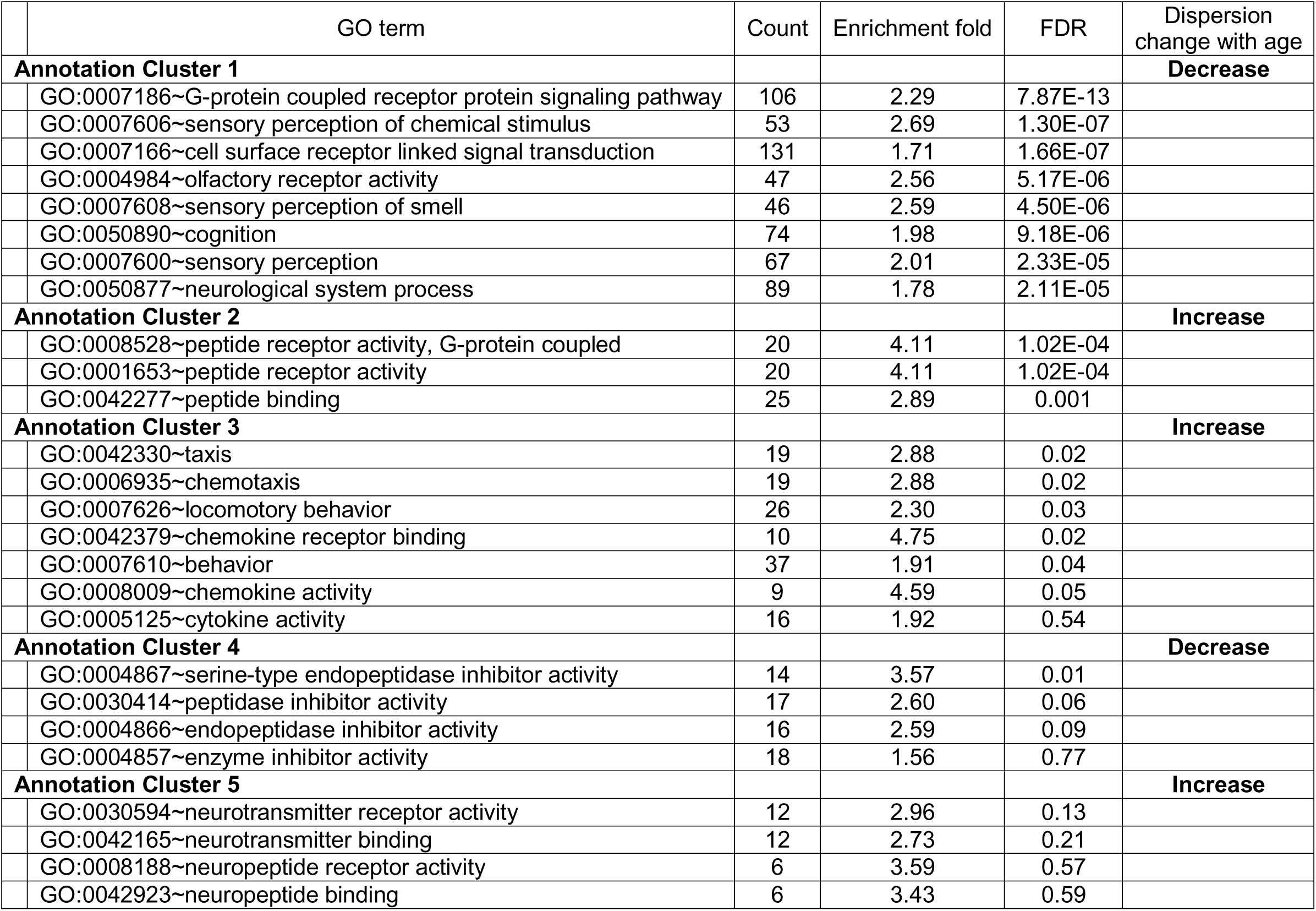
Top five GO-term clusters for DV genes showing significant differential variability in their expression between age groups.

Furthermore, we expanded the utilization of Levene’s test, coupled with a distance measure, to a multivariate setting (Anderson, 2006) to identify age-related DV gene sets, i.e., sets of genes with significant differential expression dispersion between young and old age groups (**Methods**). Eight GO term-defined gene sets (seven distinct gene contents) were identified at the 5% FDR significance level in three brain regions (**Table 3**). These include a set of two genes, *TERF1* and *TERF2*, with the function of age-dependent telomere shortening. Lin et al. (2014) showed that TERF1 and TERF2 use different mechanisms to find telomeric DNA but share a novel mechanism to search for protein partners at telomeres. The deviation of expression profiles of the two genes from individual samples to the population mean centroid was measured with Mahalanobis distance (MD) (**Methods**). Compared to old specimens, young samples show an increased level of scattering in their *TERF1-TERF2* expression, indicated by the higher level of MD (**Fig. 4C**).

**Table 3.**
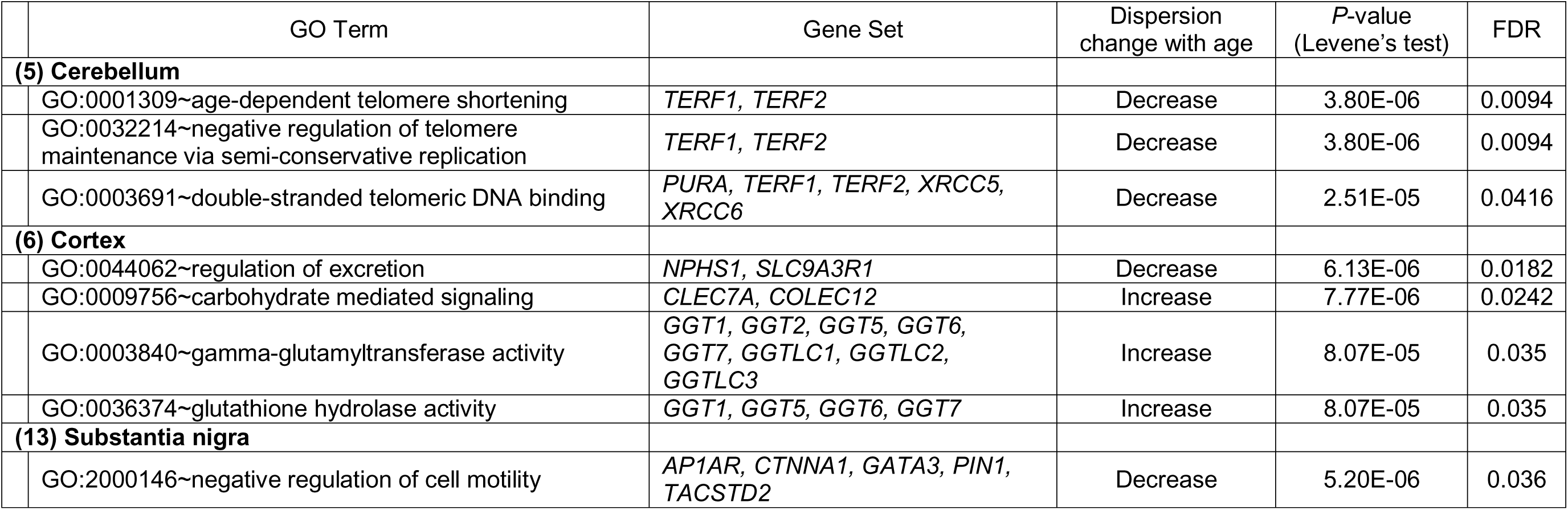
Gene sets whose gene expression dispersion differs significantly (FDR<5%) between young and old adults. Genes of each set were grouped based on the shared GO term in their functional annotations.

## Discussion

Using the GTEx data, we assessed the associations between gene expression and chronological age in different neuroanatomical regions of the human brain. The main findings of our work include: (1) the gene expression responsiveness to aging in various brain regions varies widely, and (2) the gene expression dispersion is a biologically relevant parameter for characterizing the age-related expression alteration.

Transcriptomic assays of the GTEx project generate high-dimensional structured data sets in which there are correlated patterns across large numbers of genes. Some of these are due to the known technical or biological effects, which can be removed by fitting them as covariates. However, even after this, there is typically substantial structural correlation that can potentially confound the subsequent analyses (Leek and Storey, 2007;Parts et al., 2011). Therefore, correcting hidden confounding factors along with covariates is indispensable in revealing the true relationship between gene expression change and the effect under consideration—which, in our case, is aging. Thus, we have carefully controlled for the structural correlations in different brain regions by inferring the hidden confounding factors using the method of factor analysis (Stegle et al., 2012) and then regressing them out. As a result of a rigorous control of input data, we detected a significant number of age-related genes (1,446 distinct genes at the 5% FDR level) across brain regions, which is more than what have previously reported elsewhere (Lu et al., 2004;Glass et al., 2013).

### Aging of the brain occurs in a region-specific manner

Our analysis of age-related genes specifically focused on each sub-region of the human brain. We detected the most genes with significant age associations in the cerebellum, which plays an important role in adapting and fine-tuning motor programs to make accurate movements, as well as the cortex, which plays a major role in many complex brain functions such as memory and awareness.

At this stage, we were aware of the effect of freezing—the frozen storage seemed to have a profound impact on the age-related gene detection. With the expression data from cerebellum and cortex samples subjected to frozen storage, we were unable to detect as many age-related genes as we identified using data from unfrozen "fresh" cerebellum and cortex samples. That is to say, the linear regression-based method for detecting age-related genes was unpowered when applied to the frozen brain samples.

Nevertheless, samples of the majority of brain regions (11 out of the 13) analyzed in the present study were collected from frozen brains. We considered them to be processed in a uniformly consistent manner and thus the results generated from these brain regions are comparable to each other. The relative abundance of age-related genes detected in these regions suggests that different regions have different age-related gene expression changes as a result. To the best of our knowledge, this is the first time that expression data was analyzed for so many brain regions in large numbers of samples processed similarly across locations and times for a single study. Overall, our results support the idea that in the human brain there are measurable patterns of gene expression changes associated with age, and these patterns are distinct from one region of the brain to another. Given that the effect of freezing tends to weaken the overall differential expression signal, our results of the number of age-related genes derived from frozen samples of the 11 regions should be considered as a lower bound of the real number of age-related genes.

### Pathway-based factor analysis identifies functional gene sets related to age

We adopted a newly developed pathway-based factor analysis (Anand Brown et al., 2015) to identify age-related gene sets. The analysis is a two-step approach. The factor analysis method, implemented in PEER (Stegle et al., 2012), was first used to discover patterns of common variation across the entire data set. Then newly derived factors summarizing expression of pathways or gene sets were used to analyze the relationships between expression and aging. This analysis allowed us to identify functionally related genes with a common response to aging. Our results support that aging is associated with a large number of biological processes and molecular functions. Many of these associations are consistent with our current knowledge. For example, aging is related to chromatin modulation (Feser and Tyler, 2011), apoptotic signaling pathway (Harman, 1992), glutathione and vitamin (Nuttall et al., 1998), oxidation-reduction process (Berlett and Stadtman, 1997), spliceosome complex assembly (Rodríguez et al., 2015), and neurotransmitter transport (Segovia et al., 2001).

In addition to focusing on linear patterns of gene expression change with chronological age, we also observed extensive interactions between the aging effect and the influence of background regulatory variants. These findings are important for the in-depth analysis of aging effects from the perspective of personal genomics.

### Evidence for the aging effect on gene expression dispersion

There is a substantial body of evidence for the impacts of aging on gene expression dispersion. In mice, for example, Southworth et al. (2009) observed a decrease in the correlated expression between normally co-expressed genes, which was associated with aging. Also in mice, Bahar et al. (2006) demonstrated an age-related increase in cell-to-cell gene expression variation in the heart [but see (Warren et al., 2007)]. Data from both humans and rats indicate that gene expression becomes more heterogeneous with age (Somel et al. (2006), and Li et al. (2009) further showed that gene expression variability in male rats is age-dependent. More recently, using human twin data, Oh et al. (2016) found that gene expression levels as well as epigenetic modifications increased in similarity in brain tissues of older individuals. Furthermore, a large study by Peters et al. (2015) revealed age-related gene expression levels that decreased with age; these findings could be attributable to dysregulation of transcriptional and translational systems.

In the present study, we identified a large number of genes whose population-level expression dispersion is age-related. For example, the variance in *MS4A4E* expression in the hippocampus was greatly increased in individuals over age 60. Such an increase in expression variability may have resulted from a decrease in normal regulation of cell growth and inflammation, which may be related to an increase in AD risk (Akiyama et al., 2000;Hollingworth et al., 2011;Naj et al., 2011;Heppner et al., 2015). The increased gene expression variability may also be due to the interaction between *MS4A4E* with other genes, e.g., *CLU* (Ebbert et al., 2015). Furthermore, we argue that the incomplete penetrance observed in neurodegenerative diseases (Rossor et al., 1996;Healy et al., 2008) may be attributed to differences in phenotypic robustness, which may be associated with or reflected in the age-related gene expression variability among individuals who are susceptible to these diseases.

Also, it is interesting to explore possible mechanisms underlying the increase or decrease in gene expression variability, as global gene expression is under stabilizing selection (Khaitovich et al., 2004;Lemos et al., 2005). Previously, we have shown that both common and rare genetic variants may confer regulatory function to contribute to gene expression dispersion (Hulse and Cai, 2013;Wang et al., 2014;Zeng et al., 2015). In particular, common genetic variants contribute to gene expression variability via distinct modes of action—e.g., epistasis and destabilizing mutations (Wang et al., 2015). Rare and private regulatory variants have been found to be responsible for extreme gene expression in outlier samples (Montgomery et al., 2011;Zeng et al., 2015;Zhao et al., 2016). Given this background information about the genetic regulation of gene expression, we argue that aging may be associated with gene expression through age-related genome instability. Mutations accumulate with age in a tissue-specific manner. The major components of the mutation spectrum include point mutations and genome rearrangements such as translocations and large deletions (Busuttil et al., 2007a). The accumulation of somatic mutations over time in various tissues and organs has been suggested as a general explanation of aging (Szilard, 1959;Curtis, 1963;Vijg, 2004). Different organs or tissues show greatly different rates of mutations that accumulate with age. The brain as a whole does not seem to accumulate mutations with age at all, but certain regions of the brain (e.g., hippocampus and hypothalamus) are much more susceptible to mutagenesis and do show increased mutational loads at old age (Busuttil et al., 2007b).

In conclusion, we demonstrate that age-related gene expression is brain region-specific, genotype-dependent, and both mean and dispersion changes in expression level are associated with the aging process. These findings provide a necessary foundation for more sophisticated gene expression modeling in the studies of age-related neurodegenerative diseases.

## Acknowledgements

We thank Oliver Stegle and Leopold Parts for technical help with PEER. We acknowledge the Texas A&M Institute for Genome Sciences and Society (TIGSS) for providing computing resources and system administration support. This work was supported by the fund of China Scholarship Council to JG, and the National Natural Science Foundation of China (No. 61573296), the Specialized Research Fund for the Doctoral Program of Higher Education of China (No. 20130121130004), the Fundamental Research Funds for the Central Universities in China (Xiamen University: Nos. 201412G009, 201510384106) to GJ. The Genotype-Tissue Expression (GTEx) Project was supported by the Common Fund of the Office of the Director of the National Institutes of Health. Additional funds were provided by the NCI, NHGRI, NHLBI, NIDA, NIMH, and NINDS. Donors were enrolled at Biospecimen Source Sites funded by NCI\SAIC-Frederick, Inc. (SAIC-F) subcontracts to the National Disease Research Interchange (10XS170), Roswell Park Cancer Institute (10XS171), and Science Care, Inc. (X10S172). The Laboratory, Data Analysis, and Coordinating Center (LDACC) was funded through a contract (HHSN268201000029C) to The Broad Institute, Inc. Biorepository operations were funded through an SAIC-F subcontract to Van Andel Institute (10ST1035). Additional data repository and project management were provided by SAIC-F (HHSN261200800001E). The Brain Bank was supported by a supplement to University of Miami grants DA006227 & DA033684 and to contract N01MH000028. Statistical Methods development grants were made to the University of Geneva (MH090941 & MH101814), the University of Chicago (MH090951, MH090937, MH101820, MH101825), the University of North Carolina – Chapel Hill (MH090936 & MH101819), Harvard University (MH090948), Stanford University (MH101782), Washington University St Louis (MH101810), and the University of Pennsylvania (MH101822). The data used for the analyses described in this manuscript were obtained from: the GTEx Portal on 10/29/2015 and dbGaP accession number phs000424.v6.p1 on 10/30/2015.

## Author contributions

JJC conceived and designed the study. CLB, JG, GJ and JJC conducted the analyses. CLB, JG, GJ, and JJC wrote the manuscript.

## Competing interests

The authors declare no competing interests.

